# K-PAM: A unified platform to distinguish *Klebsiella* species K- and O-antigen types, model antigen structures and identify hypervirulent strains

**DOI:** 10.1101/2020.03.21.001370

**Authors:** L Ponoop Prasad Patro, Karpagam Uma Sudhakar, Thenmalarchelvi Rathinavelan

## Abstract

A computational method has been developed to distinguish the *Klebsiella* species serotypes to aid in outbreak surveillance. A reliability score (estimated based on the accuracy of a specific K-type prediction against the dataset of 141 distinct K-types) average(ARS) that reflects the specificity between the *Klebsiella* species capsular polysaccharide biosynthesis and surface expression proteins, and their K-types has been established. ARS indicates the following order of potency in accurate serotyping: Wzx(ARS=98.5%),Wzy(ARS=97.5%),WbaP(ARS=97.2%),Wzc(ARS=96.4%),Wzb(ARS=9 4.3%),WcaJ(ARS=93.8%),Wza(ARS=79.9%) and Wzi(ARS=37.1%). Thus, Wzx, Wzy and WbaP can give more reliable K-typing compared with other proteins. A fragment-based approach has further increased the Wzi ARS from 37.1% to 80.8%. The efficacy of these 8 proteins in accurate K-typing has been confirmed by a rigorous testing and the method has been automated as K-PAM(www.iith.ac.in/K-PAM/). Testing also indicates that the use of multiple genes/proteins helps in reducing the K-type multiplicity, distinguishing the K-types that have identical K-locus(like KN3 and K35) and identifying the ancestral serotypes of *Klebsiella* spp. K-PAM has the facilities to O-type using Wzm(ARS=85.7%) and Wzt(ARS=85.7%) and identifies the hypervirulent *Klebsiella* species by the use of *rmpA,rmpA2,iucABCD,iroBCDN* and *iutA* marker genes. Yet another highlight of the server is the repository of the modeled 11 O- and 79 K - antigen 3D structures.

## Introduction

*Klebsiella* species (spp) cause a wide-range of transmissible infections [1, 2]. Two cell surface associated glycoconjugates, namely, the capsular polysaccharides (CPS) and lipopolysaccharides (LPS) are the major virulence factors of *Klebsiella* spp [3, 4]. While the O-antigens form the outermost component of LPS, the CPS or K-antigens form the bacterial capsule. *Klebsiella* strains can effectively be discriminated based on the K- (alternatively, capsular) and O-serotyping [3]. Currently, 141 K-types and 13 O-types are used in the serotyping of *Klebsiella* spp.

*Klebsiella* spp utilize Wzx/Wzy dependent pathway for the CPS biosynthesis and surface expression, which begins with the synthesis of the nucleotide sugar precursors corresponding to a particular K-type and the assembly of the repeating unit in the cytoplasmic region with the help of the sugar specific glycosyl transferases (GTs) (**Figure 1 (top)**) [5-7]. It is worth noting that the glycosylation of the repeating unit is initiated either by WbaP (when the initializing sugar linked to the undecaprenol-pyrophosphate (Und-PP) is galactose) or by WcaJ (when the initializing sugar linked to Und-PP is glucose) [6, 7]. Followed by this, flipping of the repeating unit to the periplasmic side takes place with the help of the flippase, Wzx. Subsequently, the repeating unit polymerization is facilitated by the Wzy copolymerase [8, 9]. Finally, Wza (an outermembrane translocon), Wzc (a tyrosine autokinase) and Wzb (a phosphatase) synergistically transport the CPS onto the bacterial surface. The surface expressed CPS is then anchored onto the *Klebsiella* outer membrane protein Wzi (**Figure 1 (top)**) [10, 11]. This CPS exportation pathway is common to all the *Klebsiella* spp as all the *Klebsiella* spp have the *cps* locus genes that are essential for the repeating unit assembly, surface exportation and surface anchorage (**Figure 1 (bottom)**). Nonetheless, the LPS biosynthesis and transportation to the outer membrane of *Klebsiella* spp follow a complex pathway as LPS comprises a lipidA, a core oligosaccharide and a polysaccharide unit [7, 12]. Among these, the polysaccharide region (commonly known as O-antigen) varies across the *Klebsiella* spp and is used in the serotyping. The O-antigen synthesis takes place in the cytosol in a similar way as the CPS biosynthesis through the participation of several O-antigen specific GTs. After polymerization, the O-antigens are transported to the periplasmic-leaflet of the inner membrane by an ABC transporter that comprises two transmembrane domains (TMDs) (termed as Wzm) and two nucleotide-binding domains (NBDs) (termed as Wzt) [13]. This O-antigen ABC transporter system is common to most of the Gram-negative bacteria. Intriguingly, the ABC transporter corresponding to some of the O-types (for instance, O3, O4, O5 and O12) has an additional carbohydrate-binding domain (CBD) that is fused to the C-terminus of the NBD [14-16].

**Figure 1.**
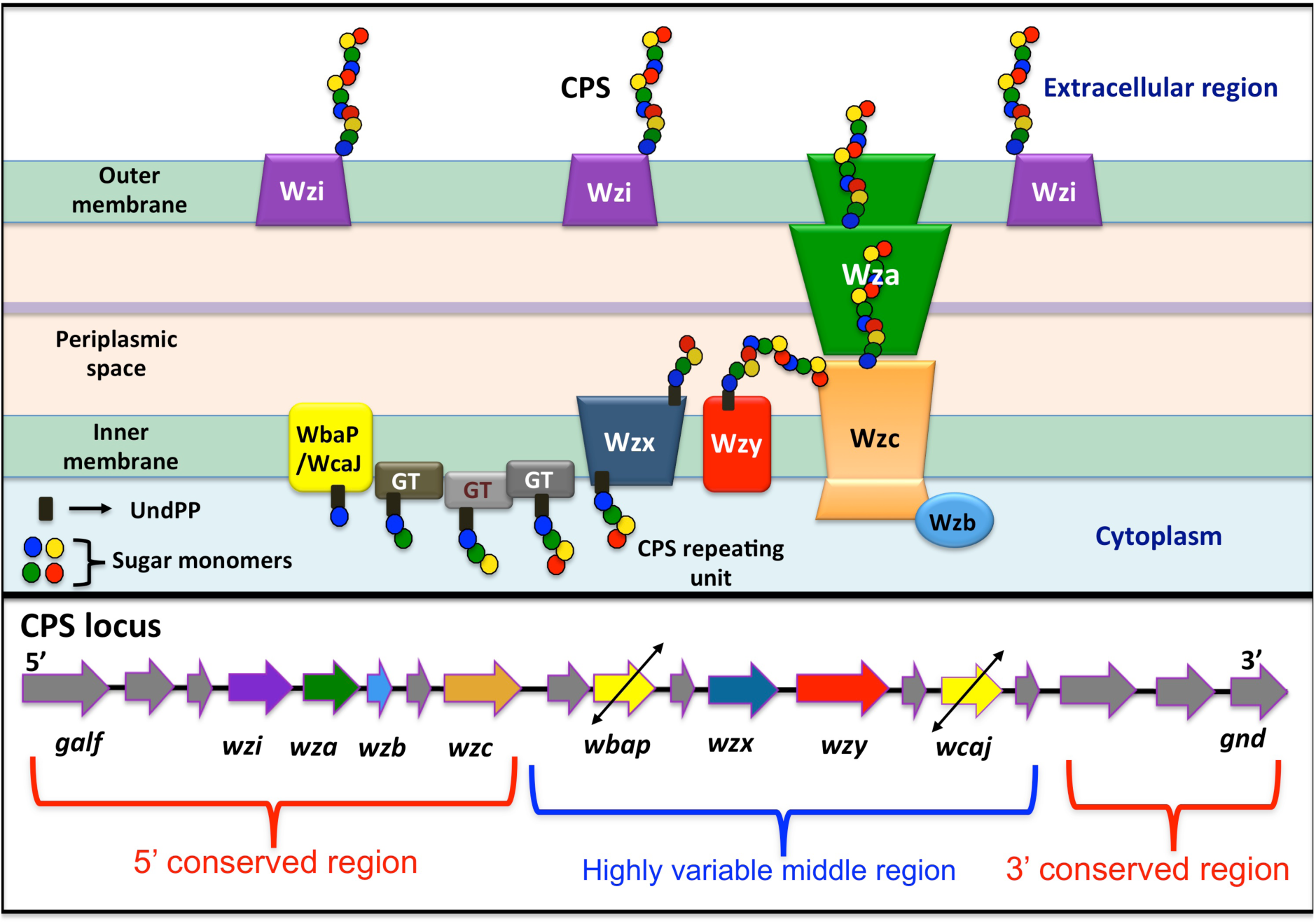
Schematic illustration of (top) the Wzx/Wzy dependent pathway involved in the *Klebsiella* spp CPS transportation and surface expression, and (bottom) the representative *cps* locus arrangement.

Capsule typing (or K-typing) using *wzi* or *wzc* genes of the CPS biosynthesis gene cluster [17] and O-typing using *wzm* or *wzt* genes involved in the O-antigen transport are quite common [18, 19]. K- & O- typing using the arrangement of genes in the corresponding biosynthesis locus is also shown to be successful [20]. Although the potential use of the other *cps* locus genes (other than *wzi* & *wzc*) in K-typing has been indicated previously [21], there is no detailed investigation on their genotype-serotype specificity and its utilization in the serotype prediction of *Klebsiella* spp. However, the genes present in the *cps* locus vary between different K-types [22] due to the variation in the sugar composition, steroisomoeric form, substituent and linkage. Luckily, *wbaP, wcaJ, wzx, wzy, wzc, wzb, wza* and *wzi* are common among all the K-types [22] as these genes are essential for the assembly, transport and surface anchorage of the K-antigen. One can envisage that among these eight genes, *wbaP, wcaJ, wzx, wzy* and *wzc* may be highly K-type specific due to their participation in the assembly of the repeating unit and its multimerization. Further, *wbaP* and *wcaJ* are mutually exclusive. Thus, K-typing with the use of these eight genes may be a reliable and time saving option, instead of considering all the *cps* locus genes (there are 508 unique protein-encoding genes) [22] for K-typing. In this context, we have classified the gene and protein sequences of *wzi, wza, wzb, wzc, wzx, wzy, wbaP* and *wcaJ* according to different *Klebsiella* spp K-types and demonstrated their importance in K-typing. A detailed analysis clearly indicates that the Wzx, Wzy and WbaP sequences are highly reliable for K-typing compared with other protein sequences. Owing to the fact that WbaP and WcaJ are mutually exclusive, Wzy and Wzx are found to be proficient K-type specific markers compared with other protein sequences mentioned above. Further, the use of multiple gene/protein sequences in reliable K-typing has been demonstrated. Similarly, O-typing of *Klebsiella* strains using *wzm* or *wzt* genes has been successfully tested.

Aforementioned approach has been implemented as an automated *Klebsiella* species K-/O-typing web server, namely, K-PAM (*Klebsiella* species serotype predictor and surface antigens modeler) (www.iith.ac.in/K-PAM/). The server accepts both the amino acid and nucleotide sequences corresponding to a single gene or multiple genes or the entire *cps* locus or the whole genome. The server also has the feature to distinguish the newly emerged hypervirulent *Klebsiella* strain (hvKp) from the classical strain (cKp) with the use of *rmpA, rmpA2, iucABCD, iroBCDN* and *iutA* genes (these genes are absent in the cKp) [23, 24]. The server also has the 3D structural repository of 75 K (K1 to K83 except K29, K42, K52, K65, K75-78) [21] and 11 O (O1, O2a, O2c, O2aeh, O2afg, O3, O4, O5, O7, O8 and O12) [25] antigens. The modeled K- and O-antigen repository may be useful in the perspective of designing polyvalent anti-*Klebsiella* vaccines [26-29] against a greater number of *Klebsiella* spp.

Thus, the K-typing methodology developed and implemented as an automated web server in the current investigation would act as a rapid and accurate *in silico* diagnostic tool for the seroepidemiological and clinical investigations through aiding in the identification and monitoring of the *Klebsiella* spp.

## Methodology

### Serotype specific classification of eight K-locus and two O-locus genes

To effectively use the eight K-locus genes of *Klebsiella* spp in K-type prediction, *wzi, wza, wzb, wzc, wzx, wzy, wbaP* and *wcaJ* gene and their protein sequences (**http://iith.ac.in/K-PAM/doc.php**) are collected from the NCBI or from the Kaptive web [20] and classified according to 141 K-types. These K-types include 77 K-antigens (*viz*., K1–K81, excluding K75–K78) of *Klebsiella* spp whose capsule types are defined serologically [21](as well as additional K-types that have been identified based on the *cps* locus or K-locus (KL) arrangement. The latter is known as the KL series (KL1–KL81, KL101–KL149, KL151, KL153–KL155, and KL157–165) [22]. It is noteworthy that the K1–K81 K-types are also synonymously referred to as KL1–KL81 locus types. However, the sugar compositions of the remaining antigens in the KL series are unknown. Similarly, the *wzm* and *wzt* genes that are involved in the transport of *Klebsiella* O-antigens are being employed in the O-typing [18, 19]. Thus, *wzm* and *wzt* gene and the corresponding protein sequences are collected and classified according to their O-type (http://iith.ac.in/K-PAM/doc.php).

### Diversity of the proteins used in K- and O-typing

The multiple sequence alignment (MSA) and the percentage identity matrix (PIM) between the protein sequences corresponding to different K-/O-types have been generated individually for all the 10 proteins using ClustalOmega [30] to derive the information about their sequence diversity among different K-/O-types. Subsequently, the region specific diversity of the ten proteins has been analyzed individually from the amino acid sequence logo generated using Weblogo3 [31].

### Reliability score

Although PIM provides the information about the degree of diversity for the eight different CPS proteins across different K-types, it may not reflect the specificity towards different K-types as it simply contains the sequence identity information. Thus, to quantify the ability of the eight K-antigen (Wza,Wzb,Wbc,Wzi,Wzx,Wzy,WbaP and WcaJ) and two O-antigen (Wzm and Wzt) biosynthesis/transport proteins in accurate serotyping (*viz.*, their antigen specificity), a reliability score (RS) has been introduced. The RS for each protein is estimated individually for 141 K-serotypes and 13 O-serotypes based on their ability in predicting a unique serotype when searched against the reference dataset using a 98% sequence identity cutoff:

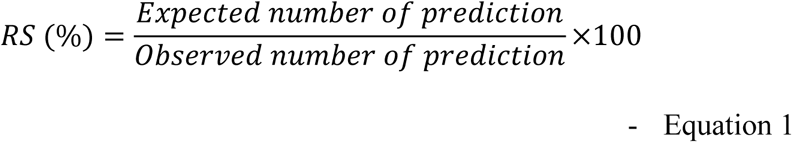

For the RS calculation, only the unique datasets (non-identical) from each K-type is retained in the reference database. Note that the cutoff is chosen to be 98% as most of the non-identical sequences corresponding to the same serotype falls above 98% sequence identity and indeed, reducing the cutoff may reflect uniformly in the RS score of all the proteins. Thus, it may not affect the reliability trend. The “expected number of prediction” (numerator in **Equation 1**) is always considered as 1 to reflect the one-to-one correspondence between the sequence and the serotype. The RS becomes 100%, when a protein sequence searched against the database gives a unique K-type. On the other hand, when 2 possible serotypes are predicted, the RS becomes 50%. Subsequently, an average reliability score (ARS) for each protein is calculated by,

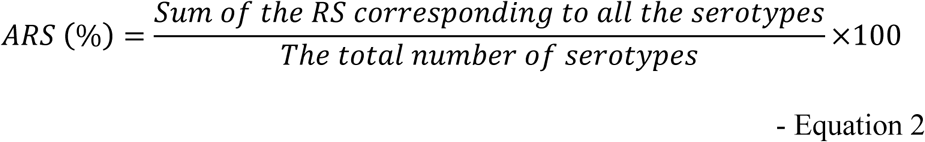

The RS and ARS calculated for different K- and O-types are given in the tables S1 (A) and S1 (B) respectively.

### Automation of serotype prediction

#### Construction of a local database

A local database comprising *Klebsiella* spp *wza, wzb, wzc, wzi, wzx, wzy, wbaP* and *wcaJ* gene as well as their protein sequences (whose K-types are already known) has been constructed by fetching the FASTA format sequences from the NCBI (except for KL104, KL107, KL128, KL145, KL147, KL151, KL152, KL155, KL157, KL158 and KL160-KL165, for which, the sequences are taken from the published literature [20]) and are grouped according to 141 distinct K-serotypes. The database consists of 1345 and 1095 non-redundant gene and protein sequences respectively corresponding to 8 genes in the *cps* locus. To perform O-type prediction, 42 *wzm* and 44 *wzt* gene sequences and 21 Wzm and 23 Wzt protein sequences that are classified according to thirteen O-types have been incorporated in the database.

A Linux based web server, namely, K-PAM (https://www.iith.ac.in/K-PAM/) that hosts K-/O-types prediction contains the database of the reference sequences that are classified according to different K-/O-types.

#### K-typing

The gene or protein sequences (FASTA format) corresponding to one or more CPS genes (*wza,wzb,wzc,wzi,wzx,wzy,wbaP* and *wcaJ*) or proteins can be used as the input for K-type prediction. Additionally, the user has the option of directly specifying the GenBank ID. As the whole genome sequencing (WGS) is becoming more popular nowadays, the server also accepts WGS sequence in FASTA or FASTQ formats from a single or multiple (contigs) files.

#### K/O-type prediction using a single gene/protein

When a single protein is given as the input, the server directly runs the BLAST [32, 33] against the local database to fetch the sequence that has 100% identity with the query. If any hit is found with 100% identity, then the K-type of the hit is assigned to the query sequence. If no reference sequence has 100% identity with the query, the server subsequently does the K-typing by filtering the hits that fall above 98% identity. If more than one hit is found, the server reports all the K-types as the possible K-types. Note that 98% cutoff is used after a close inspection of the reference dataset (https://www.iith.ac.in/K-PAM/pim.html), which indicates that, in 85% of the cases, the sequences from a same K-type have the sequence identity above 98%. However, about 15% of the sequences have the identity below 98% and above 90% within the same serotype. Thus, to improve the prediction accuracy, when no serotype is found for 98% identity cutoff, the server further reduces the cutoff sequence identity to 90% in a stepwise manner (reducing the sequence identity by 2% in each round) to fetch the appropriate serotype. If no hit is observed using the above cutoff criteria, the server doesn’t report any K-type. It is noteworthy that in a similar fashion, K-typing using the gene sequences has been performed by considering 95% and 90% as the highest and the lowest sequence identity cutoff.

#### K-type prediction using multiple genes/proteins

To make the K-type prediction more robust, an option is also given to use the multiple genes/proteins for the prediction. In this case, the server automatically does the BLAST search and identifies the genes/proteins present in the query. As described above, the K-typing is performed for the individual proteins and is reported along with a relative reliability score (calculated using **Equation 1** and see below for details). Such a combined K-typing strategy may be more promising in terms of reducing the false positive K-typing (*viz*., 100% prediction accuracy), especially in the case of less divergent Wzi and Wza sequences. When multiple K-types are predicted from different proteins, the common K-type predicted from the individual protein sequences is designated as the K-type (marked as (1) in **Figure 2**). Note that when more than one K-type is found to be common for all the proteins, the server considers both as potential K-types. If less than three proteins from the query sequence have a sequence identity above 98% with the reference dataset or if no common K-type is predicted across the multiple CPS proteins by applying 98% identity cutoff, the server reports it as a newly emerged K-type (marked as (2) in **Figure 2**). When multiple protein sequences give identical K-type (using the identity criterion of 98%) except a few, the protein sequences that give different K-type(s) are again searched against the reference database by reducing the sequence identity cutoff criterion to 95%. When no serotype is reported for 95%, the server again reduces the cutoff criterion to 90% (by taking into consideration of the sequence identity corresponding to the same serotype, which falls above 90%) in a stepwise manner as discussed above. If a common K-type is found by applying the relaxed criterion mentioned above, then, it is assigned as the variant of the K-type (marked as (3) in **Figure 2**). If no common K-type is observed even after the relaxation of sequence identity cutoff, the server reports the K-type as the variant of the K-type that is predicted from the majority of the proteins (marked as (3) in **Figure 2**). Further, when multiple K-types are found to be common in at least three different proteins, then the K-type is reported as the hybrid of both the K-types (marhed as (4) in **Figure 2**). A similar methodology has been used to K-type using the gene sequences by considering 95% and 90% as the highest and the lowest sequence (relaxed) identity cutoff.

**Figure 2.**
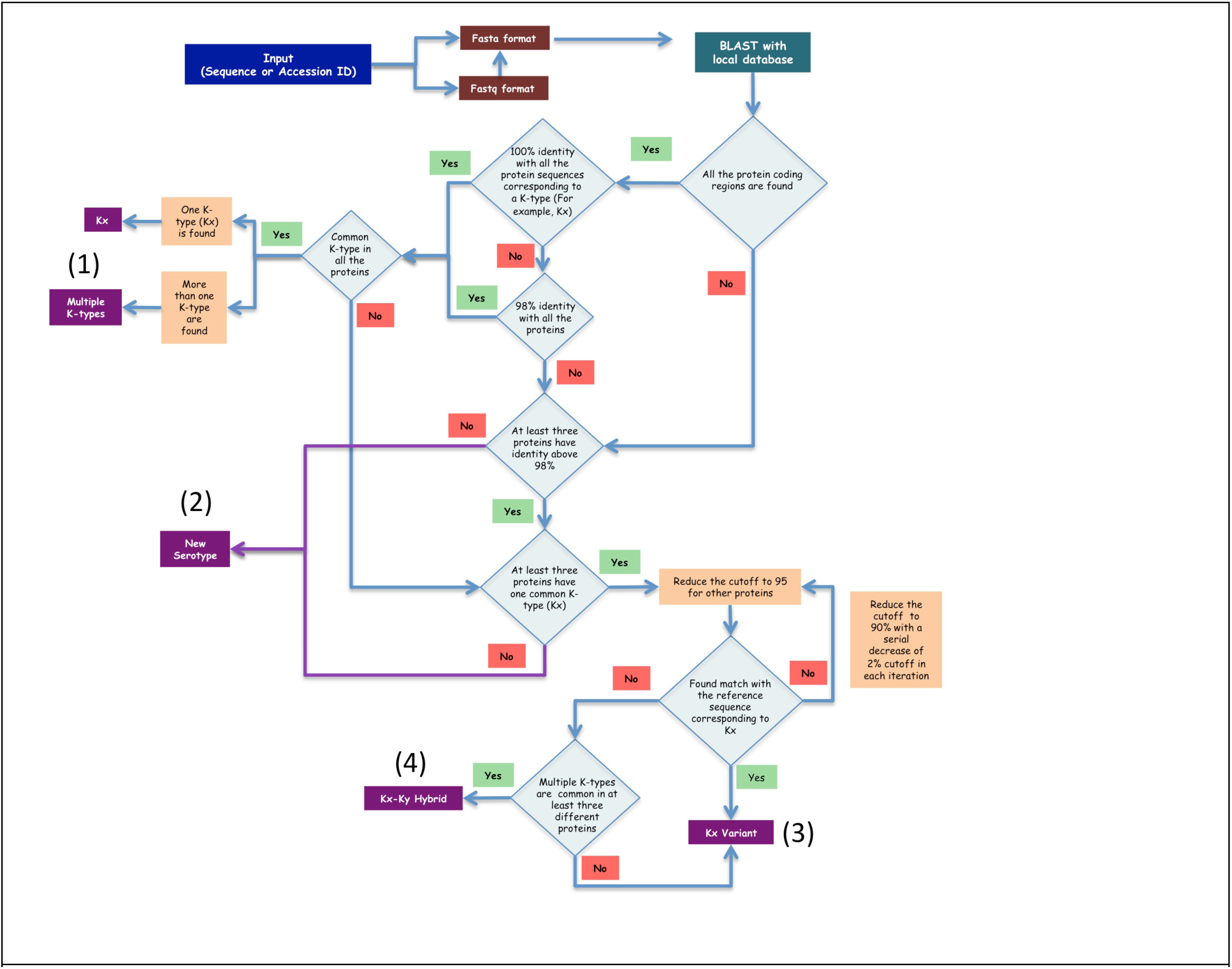
Flowchart describing the methodology implemented in K-PAM for K-typing using multiple gene/protein sequences. Note that the methodology is explained here by considering the protein sequences as example. The final serotype prediction is indicated in a violet colored box.

#### O-typing

Similar to K-typing, O-typing can be done using both gene (*wzm* and *wzt*) and protein (Wzm and Wzt) sequences. Sequence identity cutoff criteria used for K-typing are followed in O-typing also. As the Wzm and Wzt sequences of O1, O2 and O2ac are highly similar (>98%), WbbY and WbbZ sequences corresponding to O1 and O2ac have been used to distinguish them from each other as well as from O2 as suggested in the earlier studies [25, 34]. Similarly, WbdA and WbdD sequences have been added to the reference database to identify the sub-types of O3 serotype (O3/O3a & O3b) [35].

### Relative sequence identity factor and relative reliability factor

To incorporate the sequence identity percentage between the query and the reference protein sequences into the prediction weightage, a scaling factor, namely, the relative sequence identity factor (RSF) has been introduced by dividing 95%/98%=0.97 (see the section “K-type using multiple genes/proteins”). Thus, a K-type identified above 95% is given less weightage compared with a K-type, which is identified above 98% by simply multiplying the RS (estimated for each protein in an every new prediction) (**Equation 1**) of the former with 0.97. Note that the standard RS reported in **Tables S1 (A) & S1 (B)** are based on the reference dataset (consists of 1095 *cps* locus protein sequences and 45 Wzm & Wzt sequences). Similarly, to give emphasis on a protein that gives more reliable K-type compared to the other protein(s) during the multiple protein based K-typing, a relative reliability factor (RRF) (**Table 1**) has been introduced:

**Table 1.** The relative reliability factor (RRF) calculated using the ARS corresponding to the reference dataset (Table S1 (A)).

|  | **Wzx** | **Wzy** | **WbaP** | **Wzc** | **Wzb** | **WcaJ** | **Wzi** | **Wza** |
| --- | --- | --- | --- | --- | --- | --- | --- | --- |
| **Wzx** | 1 | 0.99 | 0.99 | 0.98 | 0.96 | 0.95 | 0.82 | 0.81 |
| **Wzy** |  | 1 | 1 | 0.99 | 0.97 | 0.96 | 0.83 | 0.82 |
| **WbaP** |  |  | 1 | 0.99 | 0.97 | 0.96 | 0.83 | 0.82 |
| **Wzc** |  |  |  | 1 | 0.98 | 0.97 | 0.84 | 0.83 |
| **Wzb** |  |  |  |  | 1 | 0.99 | 0.85 | 0.84 |
| **WcaJ** |  |  |  |  |  | 1 | 0.86 | 0.85 |
| **Wzi** |  |  |  |  |  |  | 1 | 0.99 |
| **Wza** |  |  |  |  |  |  |  | 1 |

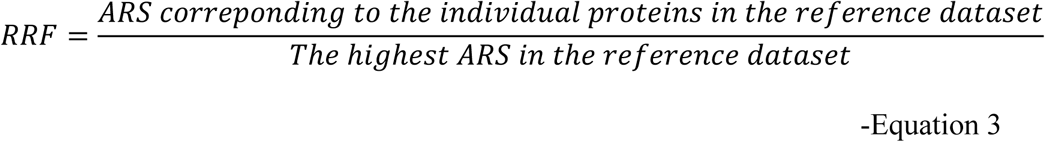

Thus, the K-type prediction from the protein that has the highest ARS (**Tables S1 (A)**) among the eight CPS marker proteins found in the query sequence is given more weightage by further multiplying the RS with the corresponding RRF (**Table 1**). Thus, both the percentage identity and the reliability are incorporated into the prediction accuracy.

### Validation of K-type and O-type prediction methodology implemented in K-PAM

The serotype prediction accuracy is validated initially by considering the sequences with known serotypes as test cases. For this, the sequences in the databases are divided into a reference dataset (consists of 141 sequences) and a test dataset (consists of 70 sequences) to corroborate the proposed algorithm. The testing against the reference dataset is done using the test dataset as a query. Further, testing is done for the sequences whose serotype is undefined in the NCBI database.

### Hypervirulent strain identification

K-PAM also has the feature to distinguish the hyper virulent *Klebsiella* spp form the classical *klebsiella* spp using the marker genes *iroB, iroC, iroD, iroN, iucA, iucB, iucC, iucD, rmpA, rmpA2* and *iutA* which are reported as hyper virulent *klebsiella* strain markers in the earlier investigations [23, 24]. It is worth noting that *iroBCDN, iucABCD* & *iutA*, and *rmpA* & *rmpA2* genes correspond to the loci of salmochelin siderophore, aerobactin siderophore and hypermucoidy respectively. These genes are specific to hypervirulent phenotype of *Klebsiella* spp, among which, *rmpA, rmpA2* and *iucABCD* are highly reliable in identifying the hypervirulent strain [23, 24]. The reference dataset of K-PAM has these gene sequences and a sequence identity cutoff criterion of 60% is used to identify the presence of these genes in the query sequence.

### Standalone version of K-PAM

A standalone version (application program interface (API)) of K-PAM is also available for analyzing the large datasets. It has the option to upload multiple files. A compressed file corresponding to Mac, Linux and Windows operating system can be downloaded from the K-PAM web server (http://www.iith.ac.in/K-PAM/kpam_doc/sk_pam.php). This standalone version has the K-typing, O-typing and hypervirulent strain identification features. The K-PAM API generates the serotype prediction summary in the CSV file format.

### Klebsiella spp K-/O-antigens 3D structural repository

In addition to the serotype prediction, the server also has the database of the modeled 3D structures of 75 K- and 11 O-antigens, the sugar composition and the glycosydic linkages within/between the repeating units are collected from the previous experimental studies [36-38] [39]. Subsequently, this information is used in the modeling of 75 K-antigen monomers using GLYCAM webserver [40]. A few unusual sugar moieties that contain substitutions like formyl group, carboxy ethyl group, cyclic pyruvate, acetyl and 4-deoxy-threo-hex-4-enopyranosyluronic have been modeled manually using Pymol [41]. These are further minimized by CHARMM (Chemistry at HARvard Macromolecular Mechanics) molecular modeling software [42] using the parameters derived from CGenFF (CHARMM General forcefield) [43]. CGenFF encompasses the parameters for a wide range of drug-like molecules, chemical groups present in the biomolecules and heterocyclic scaffolds [43]. CHARMM36 carbohydrate forcefield is used in the minimization of unmodified sugars. Minimization step includes 1500 steps of steepest descent followed by 1500 steps of adopted basis Newton-Raphson energy minimization method with a non-bonded cut-off of 16Å. During the minimization, Generalized Born with a simple switching has been used to incorporate the solvation effect implicitly [44, 45].

A total of 75 *Klebsiella* K-antigen 3D structures were modeled and a local database has been created and grouped according to *Klebsiella species* (http://iith.ac.in/K-PAM/k_antigen.html) [46]. Modeled K-antigens have been given a suffix ‘KK’ (Klebsiella K-antigen) and each sugar has been given a nomenclature depending on the substitutions and modifications as described elsewhere [47]. It is noteworthy that the 3D structures of K29, K42, K52, K65, K75, K76, K77 and K78 antigens are not modeled due to the unavailability of their sugar composition and linkage information.

In addition, the server has the repository of 11-modeled LPS structures of *Klebsiella* species [25]. The LPS structures have been modeled by simply specifying the sugar composition and linkage using CHARMM-GUI LPS modeler [48]. The modeled LPS structures consists of three parts, namely, (i) a hydrophobic lipid A (ii) core oligosaccharide linked to lipid A and (iii) an O-antigen (attached to the core oligosaccharide), a common feature of the Enterobactericeae family. The core polysaccharide is usually divided into inner and outer cores, wherein; the inner core is highly conserved in the Enterobactericeae family.

Both the O- and K- antigens can either be visualized in the JSmol web-browser or can be downloaded and viewed by any molecular visualization software.

### Multimer modeling of Klebsiella spp K-antigens

K-antigens consist of several hundred repeating units that are linked through a variety of glycosidic linkages accounting to their higher molecular weight. This makes it essential for the construction of a tool to generate K-antigen multimer. Considering this, two different multimer modeling options are available in K-PAM to generate K-antigen multimer following the methodology described elsewhere [47]: rigid multimer modeling (RMM) and flexible multimer modeling (FMM).

## Results and Discussion

### The sequence diversity of Wzi,Wza,Wzb,Wzc,Wzx,Wzy,WbaP,WcaJ,Wzm and Wzt

The sequence identity matrices reveal the following order of diversity among the proteins used for the K-typing: Wzy (the lowest sequence identity between any two serotypes is 3%) > Wzx(5%) > WcaJ(27%) > Wzb(40%) > Wzc(45%) > WbaP(61%) > Wza(65%) > Wzi(80%) (https://www.iith.ac.in/K-PAM/pim.html). The protein sequence logo corresponding to the dataset indicates that significant variation in Wzi and Wzc are confined to a particular region (**Figure S1**), whereas no such distinct region specific diversity is seen in the cases of Wza, Wzb, WbaP, WcaJ, Wzx and Wzy (https://www.iith.ac.in/K-PAM/pim.html). Similarly, the diversity is distributed throughout Wzm and Wzt sequences (https://www.iith.ac.in/K-PAM/pim.html).

### Antigen specificity quantification of proteins used for K-serotyping

The precision in the K-typing efficacy of each protein has been tested through the K-PAM web server and the results are used in the estimation of ARS (**Equations 1** and **2**). The ARS, a measure to quantify the K-type specificity, lies in the following order for CPS proteins: Wzx(ARS=98.5%) > Wzy(97.5%) > WbaP(97.2%) > Wzc(96.4%) > Wzb(94.2%) > WcaJ(93.8%) > Wza(79.9%) > Wzi(37.1%) (**Table S1 (A)**), indicating the highest prediction accuracy for Wzx as it is highly specific to each serotype (except for K22&K37 and K21&KL154). It is worth noting that K22 and K37 are the frame-shift mutants [17] and the RS estimated from **Equation 1** for K22 and K37 is 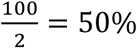. Similarly, the RS corresponding to K21&KL154 is 50% each. Wzy is mostly unique for all the serotypes, except in these cases of K22&K37, K21&KL163 and KL146&KL154 (thus, K22,K37,K21,KL163,KL146 &KL154 have the RS value of 50%). Besides these, Wzy is unique among the other serotypes. WbaP is the next highly specific protein to K-type, with the exception of K40&KL135 and KL146&KL154. It is noteworthy that the diversity in WbaP is distributed throughout the sequence (https://www.iith.ac.in/K-PAM/pim.html) and the three dimensional structural information may require in explaining the K-type specificity of the motif/domain involved in the protein’s transferase activity. As the structure of *Klebsiella* WbaP is unknown, we couldn’t pin point the K-type specific region of WbaP. Except for five pairs of K-types including K22&K37 and K9&K45, Wzc is highly specific to other K-types. Compared with Wzc, Wzb is less specific as it is identical for K1&K4 and K15&K52 in addition to K22&K37 and K9&K45. WcaJ that possesses the next ARS is identical for K16, K54, K58 & KL113 (thus the RS is 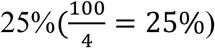) in addition to K22&K37 and K1&K4. Wza and Wzi that take the last 2 positions have many identical K-types, even Wza having 25% RS in many cases (**Table S1 (A)**). Thus, it is clear that Wzx can be the most reliable protein for K-typing. Alternatively, Wzy, WbaP and Wzc can be used for reliable K-typing. The RS for each K-type and each protein is given in **Table S1 (A)**.

### Region specific diversity of Wzi and Wzc

A careful inspection indicates that the K-type specificity is accumulated in the middle region of Wzi (falls between the two highly conserved motifs “QISAS” and “GYYQQ”) that is flanked by more conserved regions on either side (**Figure S1 (Top)**). As the periplasmic region of Wzc synergistically interacts with Wza to transport the K-antigen to the extracellular space [49], it may be highly K-antigen specific despite the sequence diversity found throughout the sequence. Thus, the periplasmic domain of Wzc can effectively be used for K-typing. Thus, a fragment based approach may be highly useful for Wzi and Wzc based K-typing. Indeed, such a fragment based approach may improve the ARS of Wzi (ARS=46%, due to the less divergence across the Wzi sequence).

### Fragment based prediction approach to improve the ARS of Wzi

In this approach, Wzi query sequence is divided into three fragments based on the marker motifs (“QISAS” and “GYYQQ”) and the K-typing is done for all the three fragments individually (**Figure S1 (top)**). To achieve precession in the prediction, the K-type assigned to all the three fragments of the query sequence is compared and the K-type that is common to all the three fragments are reported as the reliable K-type. The fragment based serotyping has significantly improved the ARS of Wzi to 80.8%, confirming the importance of fragment-based prediction approach. However, the ARS is still lower compared with all the other CPS proteins. This approach can significantly be helpful in improving the *wzi-*allele specific K-typing that is in practice [50].

Although the middle region (between the motifs “SRM” and “SVDL”) of Wzc (**Figure S1 (bottom)**) exhibits K-type specificity, rigorous testing indicates that unlike in the case of Wzi, the whole sequence of Wzc itself gives precise K-typing. Thus, to save the time on fragmentation and to speed-up the K-type prediction process, the entire Wzc sequence is used for K-typing.

### K-type prediction accuracy of K-PAM

#### Validation of the fragment based K-typing using Wzi

The K-type prediction accuracy using Wzi fragment based method (**Figure 3**, see above) has been tested by considering the sequences used for the construction of the local database. For this, all the Wzi sequences used in the reference dataset are considered as a query by removing the corresponding sequence from the database and prediction has been carried out to test the efficacy of the fragment based prediction. The results indicate that as the terminal sequences exhibit lesser diversity among different K-types, the prediction using the middle region significantly reduces the false positives (**Table S2**). The multiplicity in K-type prediction arising from the use of the N-terminal, middle and C-terminal fragments individually has been reduced by considering the K-type that is predicted commonly from the three fragments (**Figure 3)**. Nonetheless, multiple K-typing is still predicted for certain cases owing to the higher Wzi sequence identity between different K-types (**Table S1 (A)**, 80.8% ARS). Although Wzc sequence analysis indicates that the periplasmic region is highly diverged compared to the terminal regions (**Figure S1 (Bottom)**), the whole sequence of Wzc itself has a higher prediction accuracy with 98% ARS.

**Figure 3.**
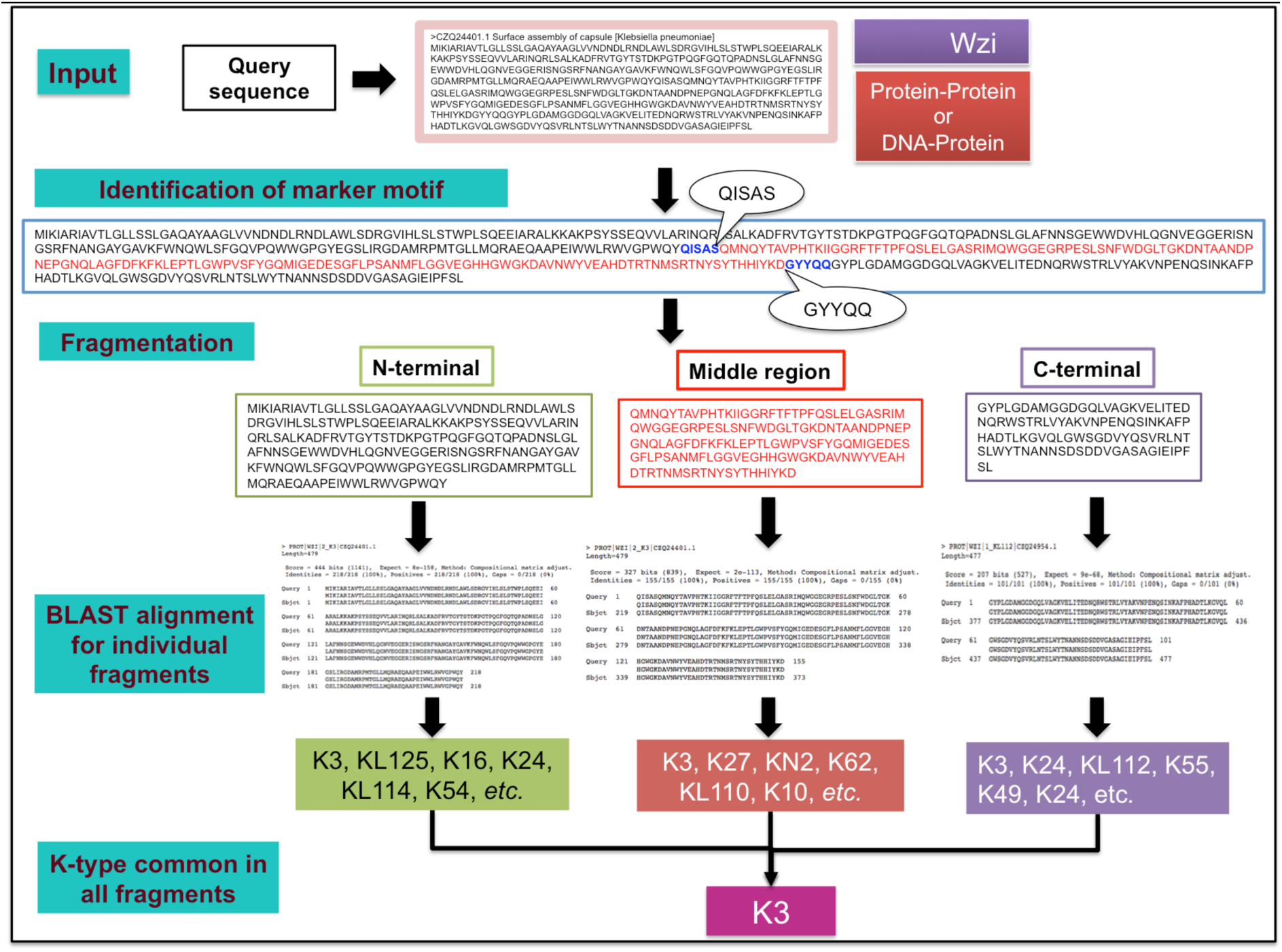
Fragment based serotype prediction method implemented for Wzi. Note that the whole query sequence gives multiple K-types (K3, KL130, KL137, K27, KL125, K62, *etc.*) for NCBI accession ID: CZQ24401.1. In contrast, three individual fragments (with respect to “QISAS” & “GYYQQ” markers) based K-typing reduces the multiplicity.

#### K-typing using multiple genes/proteins

Although *wzi*(or Wzi) or *wzc*(or Wzc) are in general used in the K-typing [17, 50], K-PAM has the option of using multiple genes/proteins for serotyping as it increases the prediction accuracy by removing the false positives. For instance, as the Wzi of K9, K38 and KL105 possess a very high sequence identity between them, when a Wzi query sequence of K38 is submitted, the server reports all the three K-types as the potential serotypes. Similarly, when the Wzc sequence of K38 alone is submitted, the server reports K38 and KN3 as predicted K-types. On the other hand, incorporation of Wza along with Wzi results in 100% prediction accuracy (http://iith.ac.in/K-PAM/doc.php). Most interestingly, K9 and K45 that have above 98% sequence identity for Wza, Wzb and Wzc can easily be distinguished on the basis of Wzx that is unique between the two (see the section **“***Serotyping of unclassified K-types***”**). Additionally, as WbaP and WcaJ are mutually exclusive genes, K9 and K45 can be discriminated based on the presence of WbaP and WcaJ respectively. Thus, using the multiple proteins increase the efficiency of K-typing (**Figure 4** and check http://iith.ac.in/K-PAM/doc.php for more such examples). Finally, a graph illustrating the reliability of prediction is displayed alongside the predicted K-type, wherein, the RS (**Equation 1**) multiplied by the relative sequence identity factor (RSF) and the relative reliability factor (RRF) (**Table 1**) for each gene/protein (X-axis) is plotted in the Y-axis (see **Methods**). The use of multiple proteins in serotyping also helps in identifying the emergence of new serotypes. For example, when different K-types are reported for different CPS genes/proteins, then, the server reports the occurrence of a hybrid (or a variant) K-type (**Figure 2**). Similarly, if the prediction from each gene/protein is random, it is reported as a new K-type. It is noteworthy that as K22 and K37 are frameshift mutants and have identical sequence identity between the 7 CPS proteins considered here, K-PAM reports the prediction with the RS of 50% even with the use of multiple proteins. A test dataset of known serotype are further considered for validating the method (**Table S3 (A)**). Except for Genbank accession ID AB819892 and AF118250, K-PAM predicts the K-types correctly. For above two cases, the K-type is predicted as the variants of K45 and K20 respectively because of the truncated query sequences. Another interesting point to be noted here is the K35 (Genbank ID: AB924573.1) and KN3 (Genbank ID: LC189075.1), whose locus arrangement is identical and the 8 gene based method of K-PAM clearly distinguishes them. This is due to the fact that the sequence identities between *wza, wzb, wzc, wzi, wzx, wzy* and *wcaJ* genes of KN3 (Genbank ID: LC189075.1) and K35 (Genbank ID: AB924573.1) are 87%, 91%, 87%, 91%, 85%, 86% and 88% respectively. Subsequently, a more rigorous testing has been carried out by considering 162 dataset from the published literature [20]. The results clearly show that K-PAM accurately predicts the serotype (**Table S4**). Further twenty-five test cases with undefined serotypes (**Table S3 (B)**) are also tested with K-PAM webserver that is discussed in detail in the later part of this manuscript.

**Figure 4.**
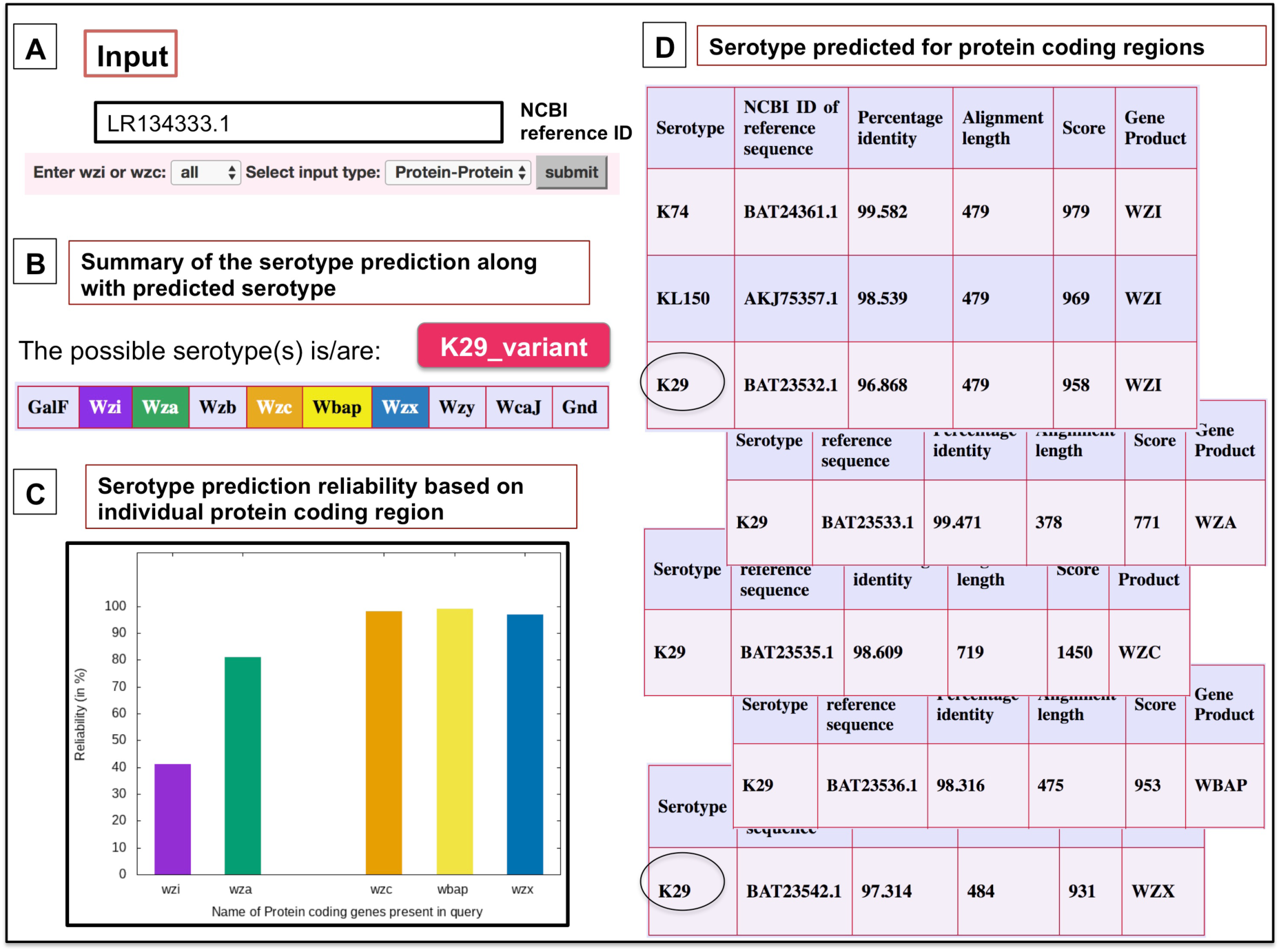
Serotype prediction example considering protein coding regions of a query sequence whose serotype is unknown (NCBI accession number LR134333.1). A) In the input box NCBI accession ID is provided along with “all” and “protein-protein” options. B) The serotype prediction summarized in the result page, wherein, the predicted serotype and coding regions identified are listed. The proteins present in the query sequence and used in serotyping are given in the color-coded box. The user can get the antigen details by clicking the serotype button. C) A graphical representation of the reliability of serotype prediction with respect to individual protein coding sequences. D) Tables (stacked) summarizing the prediction with respect to individual protein sequences. Note that after reducing the cutoff to 95%, the encircled serotype is predicted from Wzi and Wzx sequences.

Interestingly, some of the newly identified (based on the K-locus arrangements) K-types (KL101 to KL165) are found to have a sequence identity with the K1-K81. For instance, Wza, Wzb, Wzc, Wzy and WcaJ of KL104 shares 99%, 100%, 99%, 97% and 99% sequence identity respectively with K30. However, the Wzi sequence has only 96% sequence identity and Wzx sequence does not share any identity with K30. Similarly, KL106, KL135, KL142, KL148 and KL163 share close sequence identity with K22, K40, K44, K36 and K21 respectively (**Table S5**).

### The use of wzm/Wzm and wzt/Wzt in O-typing

Both Wzm (the lowest sequence identity between any two serotypes is 19%) and Wzt (the lowest sequence identity between any two serotypes is 33%) are highly divergent across the 13 O-types, thus, have become the highly suitable candidates for the prediction of O-types. Notably, the ATP binding domain of Wzt is absent in O1, O2, O8 and O9 types. Intriguingly, the ARS corresponding to Wzm (85%) and Wzt (85%) indicates that these proteins may exhibit a less O-type prediction accuracy (**Table S1 (B)**). A detailed analysis indicates that such a low value for ARS is due to the fact that O1, O2 & O2ac (RS=33%) and O3 & OL104 (RS=50%) have 100% sequence identity. However O1, O2 & O2ac and O3 variants can be distinguished with the help of the genes that are located outside the O-antigen biosynthesis locus (*rfb*) (see **Methods**).

O-type prediction using *wzt* and *wzm* gene sequences by considering the NCBI accession number CZQ25314.1 as an example has also been successfully tested (**Figure S2 (i)**).

### Application of K-PAM web server

#### Serotyping of unclassified K-and O-types

The application of K-type prediction methodology implemented in K-PAM has been demonstrated by considering several single and multiple protein sequences of *Klebsiella* species (whose serotypes are undefined) taken from NCBI **(Table S3 (B)** (multiple proteins) and check http://iith.ac.in/K-PAM/doc.php for single protein based predictions). The importance of using Wzi and Wzc sequences together has been demonstrated by considering the NCBI accession numbers CZQ24079.1 (Wzi), BAI43775.1 (Wzi), CZQ24082.1 (Wzc) and BAI43778.1 (Wzc) (**http://iith.ac.in/K-PAM/doc.php**). For instance, the multiple K-types (K15, K38, K51 and K52), which are predicted from Wzi (CZQ24079.1) alone has been narrowed down to K15 with the inclusion of Wzc (CZQ24082.1). Similarly, the K-types predicted individually for BAI43775.1 (Wzi) are K9 & KL105 and BAI43778.1(Wzc) are K9 and K45. Thus, K9 is predicted to be the K-type as it occurs in both the cases. Interestingly, K-PAM predicts K11 as the serotype for Genbank accession ID LT603705 with 100% reliability from multiple genes that was wrongly annotated as KL129 earlier [51] (**Table S3 (B)**).

Further, for Genbank ID LR134333.1 (**Figure 4**), except Wzi and Wzx all the other genes have 98% sequence identity with the database sequences corresponding to K29 (except Wzy as it is not annotated or identified in any serotypically defined K29 sequences (**Table S3 (B)**)). Further, no Wzb sequence was found in the query sequence. In this case, K-PAM relaxes the sequence identity to 95% for Wzi and Wzx to look for any match with the reference sequences corresponding to K29. It is found that Wzi and Wzx have 97% sequence identity with K29 reference sequences. Thus, the serotype is reported as a variant of K29. This reflects in the final reliability score as RS of Wzi is multiplied by RSF as well as RRF. Supplementary Table S3 (B) provides the summary of 25 test cases considered in the current investigation to illustrate the use of multiple genes/proteins in K-typing. These examples support the importance of the K-typing methodology implemented in K-PAM.

Although the server is capable of handling the whole genome sequencing that is becoming very popular and efficient, it can be useful in the serotyping of several unclassified *Klebsiella* strains whose CPS gene(s) information derived using PCR technique (wherein, only limited gene sequences are available) is accumulated in the NCBI database [52]. Thus, K-PAM will be useful in improving the serotype epidemiology and pathophysiological insights about the *Klebsiella* spp.

O-typing using Wzm and Wzt has also been predicted and validated as described above by considering the NCBI accession ID CZQ25314.1 as a case in point (**Figure S2 (i)**). Both Wzm and Wzt predict O3 as O-type. It is noteworthy that both Wzm and Wzt facilitate the prediction of O-type with 100% accuracy with the exception of O1, O2 and O2ac, and O3 and OL104 due to the high sequence identity (**Table S6**). In fact, the O-antigen biosynthesis locus arrangement is also identical for these serotypes. Thus, *wbbY* and *wbbZ* genes that are located outside the O-antigen biosynthesis locus are additionally used to distinguish O1, O2 and O2ac serotypes [25, 34]. Similarly, *wbdA* and *wbdD* genes are used to distinguish O3 variants.

### Identifying the emergence of a new serotype

K-typing methodology implemented in K-PAM is capable of not only identifying the ancestral K-type(s) that has emerged through cross-reaction, but also, a totally new K-type. For instance, the sequence identity of Wzc, Wzy, and WbaP proteins fall below 95% for Genbank ID CP029128.1, thus, the K-type predicted from them is not considered. Nonetheless, Wzi has 99.3% sequence identity with K74 and Wza has the sequence identity of 95.8% with both KL127 and KL131. Thus, K-PAM reports CP029128.1 as a new serotype (**http://iith.ac.in/K-PAM/doc.php**). Another example in this category is Genbank ID: CP035214.1.

When a particular K-type is reported dominantly from multiple (at least 3) protein sequences except one or two, K-PAM reports the new serotype as a variant of the dominant K-type. For the Genbank ID CP020358.1, the nearest K-type predicted from all the 7 proteins (Wzi, Wza, Wzb, Wzc, Wzx, Wzy and WbaP) is K29, but, the sequence identity between the reference and query sequences of Wzi is very poor. In this case, K-PAM predicts the serotype as the variant of K29. Interestingly, for the Genbank ID NZ_AP014950.1, Wzx,Wzy,Wzc,Wzb and WcaJ predict the serotype as K67. However, Wzx that has a higher ARS compared with the other proteins has only 93% sequence identity between the query and reference sequences. Thus, serotype is predicted here as the variant of K67. In addition, Wzi and Wza in the query sequence have less sequence identity (92% and 93% respectively) with K67 reference protein sequences. Thus, the methodology implemented here will also be useful in identifying the newly emerging K- or O-types.

Although *in silico* K-typing tools such as BIGSdb [53] and Kaptive web [20] are available for *Klebsiella* spp, they are either based on the arrangement of K-antigen gene cluster or based on a single gene sequence. Further, as K-PAM predicts the K-type based on multiple gene/protein sequences as well as from fragment based method, it can easily identify the antigen variants and their ancestor(s). Importantly, K-PAM precisely distinguishes the K-types that have identical CPS locus arrangement (for example, K35 and KN3).

### Hypervirulent Klebsiella strain identification

The efficacy of K-PAM in identifying the hypervirulent *Klebsiella* strains has been demonstrated by considering several clinically important strains [54-58]. **Figure S2 (ii)** depicts the hyperviruelent strain identification process of K-PAM by considering a clinically important hypervirulent *Klebsiella variicola* strain as an example [57]. Results indicate that the strain has all the hypervirulent marker genes (*iroB, iroC, iroD, iroN, iucA, iucB, iucC, iucD, rmpA, rmpA2* and *iutA*). **Table S7** summarizes the hypervirulent maker genes identified and the K-/O-types predicted by the server for the test cases considered here. It is noteworthy the server reports the strain that lacks the hypervirulent marker genes (a negative control, Genbank ID: CP022691.2) as a classical strain.

### Database of the modeled 3D structures of K-/O- antigens

A database that contains the modeled 3D-structures of the K-antigens is also incorporated in the server and upon clicking a K-antigen ID (**http://iith.ac.in/K-PAM/k_antigen.html)**, the database redirects the user to a webpage that comprises the chemical and schematic representations of the *Klebsiella* spp CPS repeating unit. The page also contains interactive Jsmol applet for the visualization of 3D structure of the K-antigen along with the provision to download the coordinates in protein databank format. The polymeric form of the K-antigens can be generated by choosing either rigid multimer modeling (RMM, **Figure S3 (**A)) or flexible multimer modeling (FMM, **Figure S3 (**B)) as described earlier in the *E coli* K-antigen 3-dimensional structural database [47]. This structural repository is user friendly as all the K- and O-antigen models are organized properly with their corresponding details and it is expected to enrich the current libraries of bacterial antigen structural libraries.

Although 2 major K-antigen classifications can be derived based on the presence of initializing galactose and glucose transferases WbaP and WcaJ respectively [7], there is no one-to-one relationship between the sequence identity of CPS proteins and the sugar compositions (**http://iith.ac.in/K-PAM/doc.php**)[7].

### Generating Klebsiella spp O-antigen models

LPS models that correspond to O-antigens, namely, O1, O2, O2aeh, O3, O4, O5, O7, O8 and O12 (modeled using CHARMM-GUI [59]) that has been deposited in the repository can be accessed through O-antigen menu bar. The LPS structures can either be visualized in the Jsmol applet or be downloaded.

## Conclusions

A detailed analysis has been carried out to test the efficacy of eight *Klebsiella* spp CPS genes/proteins namely, Wzi, Wza, Wzb, Wzc, WbaP, WcaJ, Wzx and Wzy, (which are essential for the K-antigen transportation and surface expression) in K-typing. The result indicates that WbaP/WcaJ, Wzx, Wzy and Wzc can be effectively used for K-typing due to their higher K-type specificity compared with Wzb, Wza and Wzi. The use of multiple genes/proteins individually for the K-typing reduces the K-type multiplicity. Although *cps* locus organization has become popular in K-typing due to the distinct *cps* locus arrangement between the different K-types [22], the missing information about one or more genes may mislead the K-typing. Further, the *cps* locus arrangement based K-typing may have a limitation when two K-types have an identical locus arrangement, but, have two different K-antigen repeating units, as seen in KN3 and K35. Thus, eight CPS gene/protein based K-typing approach developed, implemented (as K-PAM web server) and tested here would facilitate the accurate K-typing. The robustness of the multiple genes/proteins based K-typing approach in reducing the false-positives, reporting a variant K-type and identifying the emergence of a new K-type has been illustrated by considering several clinically important sequences. K-PAM can extensively be used to extract the K-type of the untyped *Klebsiella* strains deposited in NCBI to aid in the betterment of seroepidemiological knowledge. To our knowledge, this the first study that has precisely investigated the serotype-genotype relationship using only 8 *cps* locus genes (essential for CPS biosynthesis, transportation and surface anchorage irrespective of the serotype) in any bacterial species that use Wzx/Wzy-dependent pathway. Thus, this method can be extended to other bacterial species for rapid and accurate K-typing. Similarly, Wzm and Wzt facilitate the prediction of O-type with 100% accuracy. The server also hosts an additional feature that distinguishes the hypervirulent *Klebsiella* strain from the classical strain. Further, the server has the capability to accept the whole genome sequences in a single file or multiple files (contigs). The standalone version of K-PAM has an additional feature of handling the multiple query sequences (as multiple input files) in one go. Thus, K-PAM would be a useful diagnostic tool in the seroepidemiological and pathophysiological investigations of *Klebsiella* spp infections. The 3D repository of the modeled K-/O-antigen structures which are accessible through online may be useful in the modeling studies that require a good starting model of *Klebsiella* K-/O-antigen(s) to facilitate the design of anti-*Klebsiella* vaccines.

## List of abbreviations

CPS: capsular polysaccharide
LPS: lipopolysaccharides
RS: reliability score
ARS: average reliability score
RSF: relative sequence identity factor
RRF: relative reliability factor
GT: glycosyl transferase
hvKp: hypervirulent *Klebsiella* species
cKp: classical *Klebsiella* species
NN: nucleotide sequence searched against nucleotide sequence in the reference database
PP: protein sequence searched against protein sequence reference database
NP: nucleotide sequence searched against protein sequence in the reference database
CFG: Consortium for functional glycomics
RMM: rigid multimer modeling
FMM: flexible multimer modelling
CHARMM: Chemistry at HARvard Macromolecular Mechanics
CGenFF: CHARMM General forcefield
KK: *Klebsiella* K-antigen

## Declaration

### Ethics approval and consent to participate

Not applicable

### Consent for publication

Not applicable

### Availability of data and materials

The sequence data has been fetched from NCBI (https://www.ncbi.nlm.nih.gov/) and Kaptive reference data set (https://github.com/katholt/Kaptive/tree/master/reference_database). All the data generated or analyzed during this study has been included in this article [and its supplementary information files].

### Author Contributions

TR designed the project. LPPP and KUS executed the project. TR, LPPP and KUS wrote the manuscript.

## Acknowledgement

The authors thank IITH and CDAC for the computational resources. LPPP and KUS thank the Ministry of Human Resource Development, Government of India for the fellowships.

