## Supplementary Figures for "K-PAM: A unified platform to distinguish *Klebsiella* species K- and O-antigen types, model antigen structures and identify hypervirulent strains"

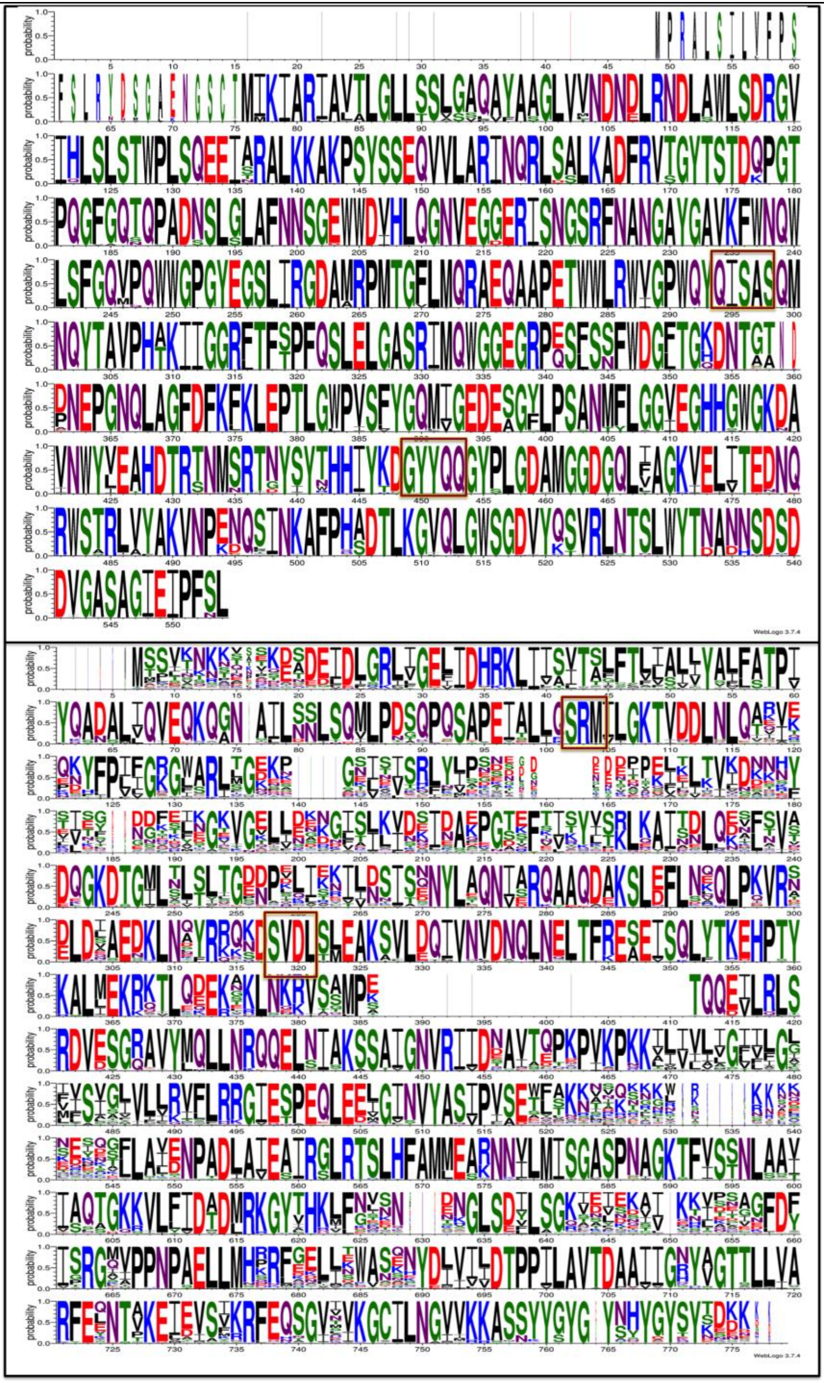

**Figure S1.** The amino acids sequence logo built using the multiple sequence alignment of 157 Wzi (**Top**) and 163 Wzc (**Bottom**) non-redundant protein sequences that are used in the creation of local database. "QISAS" & "GYYQQ" marker motifs used in the fragmentation of Wzi sequences (N-terminal, middle and C-terminal regions) and "SRM" & "SVDL" marker motifs used in the fragmentation of Wzc sequences are boxed. Note that the region falls between "SRM" & "SVDL" in Wzc corresponds to the periplasmic region of the protein that interacts with Wza to transport the K-antigens.

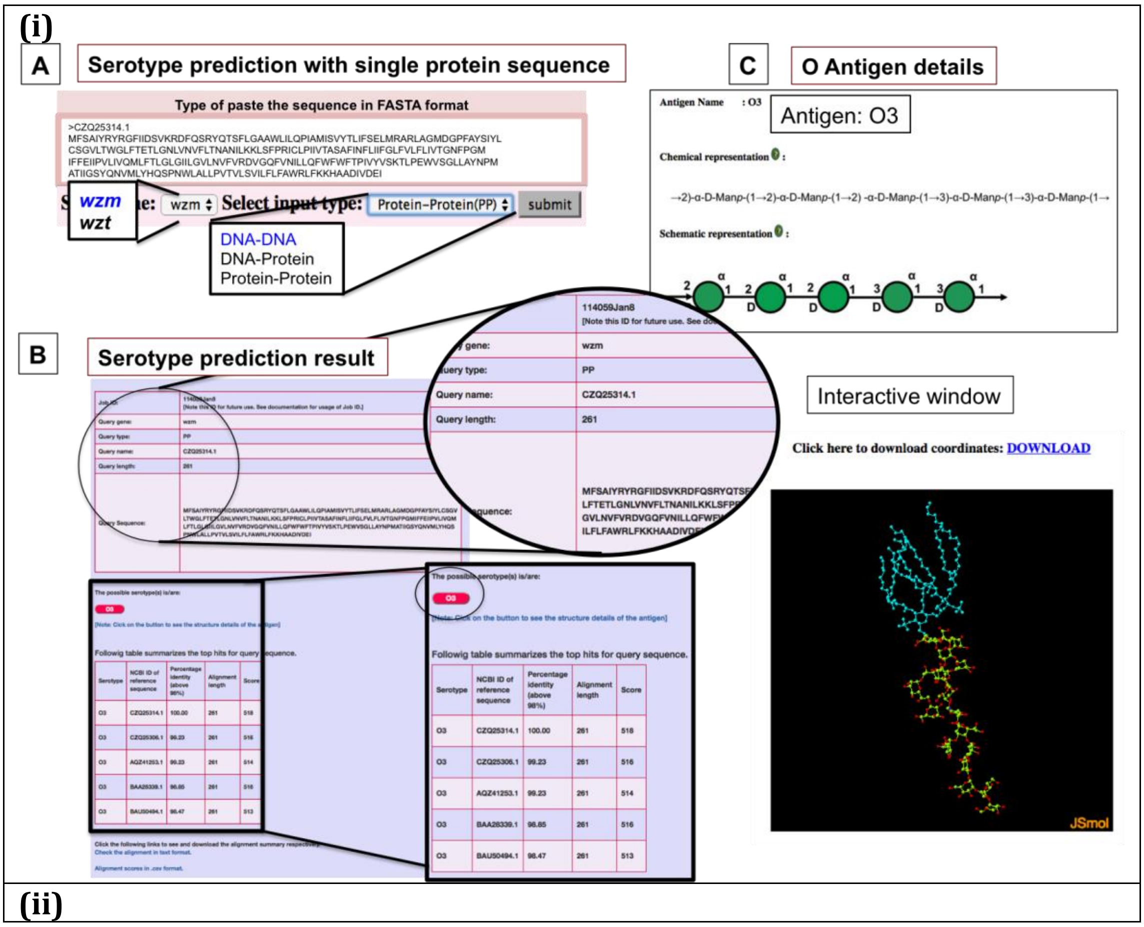
(i)
A
Serotype prediction with single protein sequence
Type of paste the sequence in FASTA format
wzm
wzt
Select input type:
Protein-Protein(PP)
submit
DNA-DNA
DNA-Protein
Protein-Protein
B
Serotype prediction result
114059Jan8
[Note this ID for future use.
Query type:
PP
Query name:
CZQ25314.1
Query length:
261
Followig table summarizes the top hits for query sequence.
C
O Antigen details
Antigen Name
Antigen: O3
Chemical representation
→2)-α-D-Manp-(1→2)-α-D-Manp-(1→2) -α-D-Manp-(1→3)-α-D-Manp-(1→3)-α-D-Manp-(1→
Schematic representation
Interactive window
Click here to download coordinates: DOWNLOAD
JSmol
(ii)

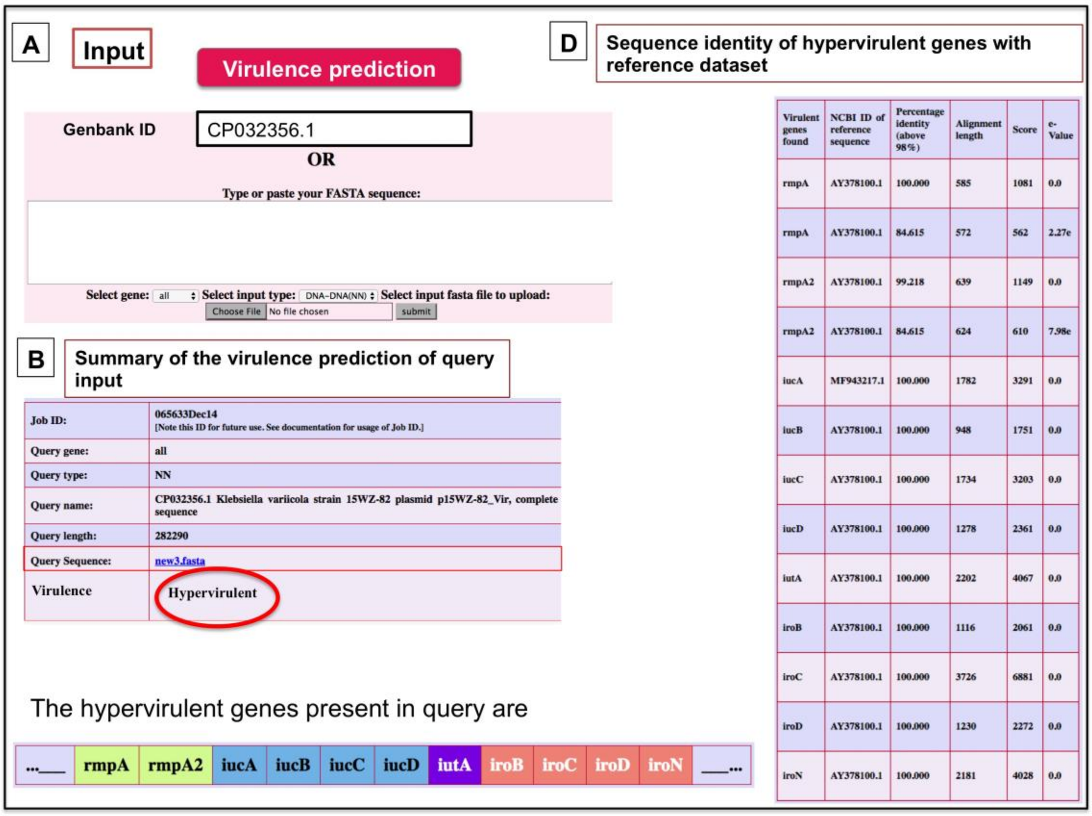


**Figure S2.** The graphical user interface of K-PAM explaining (i) the O-type prediction and (ii) the hypervirulent strain identification. **(i)** The example shown here illustrates the *Klebsiella* O-type prediction using the NCBI accession number CZQ25314.1 that correspond to Wzm. A) The input box, in which, Wzm query sequence is submitted. The options, “Wzm” and “Protein-Protein” are selected. B) O-type prediction by K-PAM is summarized in the result page that contains the reference job ID, input sequence information and the predicted O-type. The clickable button (magenta button, circled) will lead to the detailed information about the antigen structure. C) The schematic and chemical representations of the predicted antigen along with the JSmol interactive view. **(ii)** An example illustrating the hypervirlent strain identification process followed in K-PAM by considering the hypervirulent *Klebsiella variicola* clinical strain as a test case (Genbank ID CP032356.1).

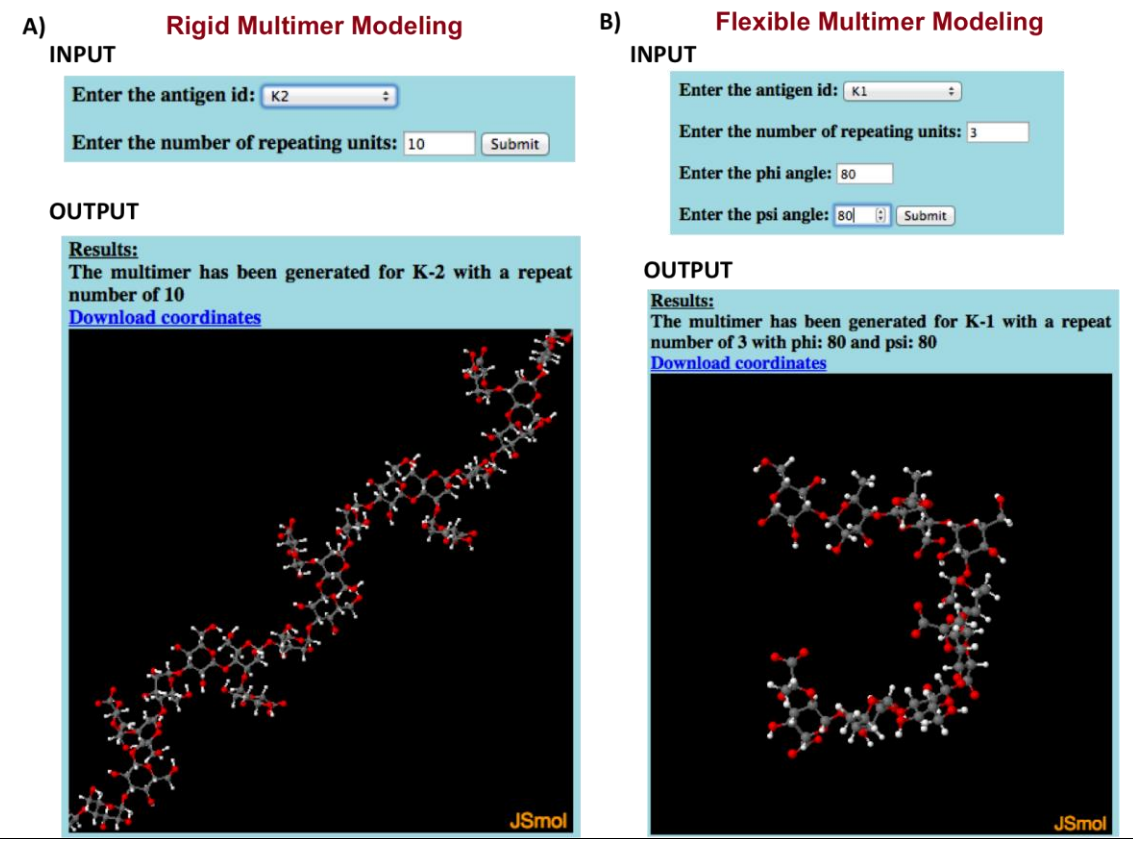


**Figure S3.** K-antigen multimer generation options. (A) Rigid multimer modeling (RMM) and (B) flexible multimer modeling (FMM) that help the user to generate the K-antigen multimers.
