## Supplementary Table S1 for "K-PAM: A unified platform to distinguish *Klebsiella* species K- and O-antigen types, model antigen structures and identify hypervirulent strains"

**Table S1.** The serotype prediction reliability calculated for (A) CPS locus proteins (for K-type) and, (B) Wzm and Wzt proteins (for O-type) by using the **Equation 1** (see the "Methodology" section in the main text). The average reliability score calculated by using the **Equation 2** (see the "Methodology" section in the main text) is given in the bottom row. **(A)** The reliability scores (column 3$^{rd}$ to 11$^{th}$ in (A) and) are color-coded as represented in the scale below. "NA" in cells stands for not applicable as, WbaP and WcaJ are mutually exclusive (either one of them is present). "NF" is mentioned for the cases where the particular protein/gene sequences for the corresponding K-type are not available. The column with title "Wzi_fbp" correspond to the reliability scores obtained after applying the fragment based approach for Wzi sequences (see the Results and Discussion section in the main text). **(B)** The reliability score (column 4$^{th}$ to 5$^{th}$) are color-coded as represented in the scale below.

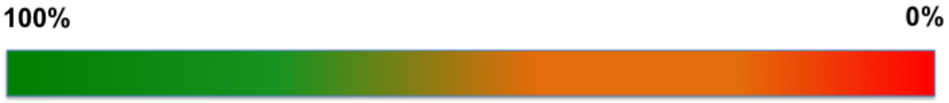


| K-type | Genbank ID or Reference ID | Reliability score in % | | | | | | | | |
|---|---|---|---|---|---|---|---|---|---|---|
| | | Wzi | Wzi_fbp | Wza | Wzb | Wzc | WbaP | Wzx | Wzy | WcaJ |
| K1 | LT174541 | 50 | 100 | 100 | 50 | 100 | NA | 100 | 100 | 50 |
| K2 | AB371296 | 25 | 100 | 25 | 100 | 100 | NA | 100 | 100 | 100 |
| K3 | LT174553 | 5.55 | 100 | 100 | 100 | 100 | 100 | 100 | 100 | NA |
| K4 | AB924548 | 50 | 50 | 33.33 | 50 | 100 | NA | 100 | 100 | 50 |
| K5 | AB289645 | 100 | 100 | 100 | 100 | 100 | NA | 100 | 100 | 100 |
| K6 | AB924549 | 50 | 100 | 100 | 100 | 100 | NA | 100 | 100 | 100 |
| K7 | AB924550 | 5.88 | 50 | 100 | 100 | 100 | NA | 100 | 100 | 100 |
| K8 | AB924551 | 14.28 | 50 | 100 | 100 | 100 | NA | 100 | 100 | 100 |
| K9 | AB371293 | 2.94 | 100 | 50 | 50 | 50 | 100 | 100 | 100 | NA |
| K10 | AB924552 | 5.55 | 100 | 100 | 100 | 100 | 100 | 100 | 100 | NA |
| K11 | LT174533 | 50 | 100 | 50 | 100 | 100 | NA | 100 | 100 | 100 |
| K12 | AB924554 | 3.22 | 33.33 | 100 | 100 | 100 | 100 | 100 | 100 | NA |
| K13 | AB924555 | 100 | 100 | 33.33 | 100 | 100 | NA | 100 | 100 | 100 |
| K14 | AB371294 | 50 | 100 | 33.33 | 100 | 100 | NA | 100 | 100 | 100 |
| K15 | LT174536 | 4.34 | 20 | 25 | 50 | 100 | 100 | 100 | 100 | NA |
| K16 | AB742228 | 50 | 100 | 100 | 100 | 100 | NA | 100 | 100 | 100 |
| K17 | AB924557 | 20 | 100 | 100 | 100 | 100 | NA | 100 | 100 | 100 |
| K18 | AB924558 | 3.44 | 33.33 | 100 | 100 | 100 | 100 | 100 | 100 | NA |

| | | | | | | | | | | |
|---|---|---|---|---|---|---|---|---|---|---|
| K19 | AB924559 | 5.26 | 33.33 | 100 | 100 | 100 | 100 | 100 | 100 | NA |
| K20 | AB289648 | 5.88 | 100 | 100 | 100 | 100 | 100 | 100 | 100 | NA |
| K21 | AB924560 | 4.76 | 100 | 50 | 50 | 50 | 100 | 50 | 50 | NA |
| K22 | AB819893 | 16.66 | 50 | 25 | 50 | 50 | NA | 50 | 50 | 50 |
| K23 | AB742229 | 50 | 50 | 25 | 100 | 100 | NA | 100 | 100 | 100 |
| K24 | AB924562 | 3.12 | 50 | 25 | 100 | 100 | NA | 100 | 100 | 100 |
| K25 | AB924563 | 14.28 | 100 | 100 | 100 | 100 | NA | 100 | 100 | 100 |
| K26 | AB924564 | 33.33 | 50 | 100 | 100 | 100 | 100 | 100 | 100 | NA |
| K27 | AB924565 | 3.84 | 33.33 | 100 | 100 | 100 | 100 | 100 | 100 | NA |
| K28 | AB924566 | 14.28 | 100 | 100 | 100 | 100 | NA | 100 | 100 | 100 |
| K29 | AB924567 | 33.33 | 50 | 100 | 100 | 100 | 100 | 100 | NA | NA |
| K30 | AB924568 | 16.66 | 50 | 50 | 50 | 50 | NA | 100 | 100 | 100 |
| K31 | AB924569 | 16.66 | 100 | 100 | 100 | 100 | NA | 100 | 100 | 100 |
| K32 | AB924570 | 100 | 100 | 100 | 100 | 100 | 100 | 100 | 100 | NA |
| K33 | AB924571 | NA | NA | 100 | 100 | 100 | NA | 100 | 100 | 100 |
| K34 | AB924572 | 100 | 100 | 100 | 100 | 100 | NA | NA | 100 | 100 |
| K35 | AB924573 | 100 | 100 | 100 | 100 | 100 | NA | 100 | 100 | 100 |
| K36 | AB924574 | 100 | 100 | 100 | 100 | 100 | 100 | 100 | 100 | NA |
| K37 | AB819894 | 14.28 | 50 | 25 | 50 | 50 | NA | 50 | 50 | 50 |
| K38 | AB924576 | 5.55 | 100 | 50 | 100 | 100 | 100 | 100 | 100 | NA |
| K39 | LT174552 | 5.26 | 100 | 20 | 100 | 100 | NA | 100 | 100 | 100 |
| K40 | AB924577 | NA | NA | 100 | 50 | 100 | 50 | 100 | 100 | NA |
| K41 | AB924578 | 100 | 100 | 100 | 100 | 100 | 100 | 100 | 100 | NA |
| K42 | AB924579 | 100 | 100 | 100 | 100 | 100 | 100 | 100 | 100 | NA |
| K43 | AB924580 | 5 | 50 | 100 | 100 | 100 | 100 | 100 | 100 | NA |
| K44 | AB924581 | 100 | 100 | 100 | 100 | 100 | NA | 100 | 100 | 100 |
| K45 | AB924582 | 100 | 100 | 25 | 50 | 50 | NA | 100 | 100 | 100 |
| K46 | AB924583 | 5 | 100 | 33.33 | 100 | 100 | 100 | 100 | 100 | NA |
| K47 | AB924584 | 3.57 | 50 | 33.33 | 100 | 100 | 100 | 100 | 100 | NA |
| K48 | AB924585 | 100 | 100 | 100 | 100 | 100 | NA | 100 | 100 | 100 |
| K49 | AB924586 | 2.7 | 100 | 100 | 100 | 100 | 100 | 100 | 100 | NA |
| K50 | AB924587 | 3.44 | 50 | 100 | NA | NA | 100 | NA | NA | NA |
| K51 | AB924588 | 3.84 | 20 | 25 | 100 | 100 | 100 | 100 | 100 | NA |
| K52 | AB924589 | 3.84 | 20 | 25 | 50 | 100 | 100 | 100 | 100 | NA |
| K53 | AB924590 | 4.76 | 100 | 100 | 100 | 100 | 100 | 100 | 100 | NA |
| K54 | AB924591 | 100 | 100 | 100 | 100 | 100 | NA | 100 | 100 | 33.33 |
| K55 | AB924592 | 4 | 100 | 100 | 100 | 100 | NA | 100 | 100 | 100 |
| K56 | AB924593 | 33.33 | 100 | 50 | 100 | 100 | 100 | 100 | 100 | NA |
| K57 | AB334776 | 3.84 | 33.33 | 33.33 | 100 | 100 | 100 | 100 | 100 | NA |
| K58 | AB924595 | 50 | 100 | 100 | 100 | 100 | NA | 100 | 100 | 50 |
| K59 | AB924596 | 100 | 100 | 100 | 100 | 100 | NA | 100 | 100 | 100 |
| K60 | AB924597 | 3.22 | 50 | 100 | 100 | 100 | NA | 100 | 100 | 100 |

| | | | | | | | | | | |
|---|---|---|---|---|---|---|---|---|---|---|
| K61 | AB924598 | 3.7 | 33.33 | 100 | 100 | 100 | NA | 100 | 100 | 100 |
| K62 | AB371295 | 4 | 50 | 100 | 100 | 100 | 100 | 100 | 100 | NA |
| K63 | AB924599 | 3.22 | 25 | 33.33 | 100 | 100 | 100 | 100 | 100 | NA |
| K64 | AB924600 | 50 | 100 | 100 | 100 | 100 | NA | 100 | 100 | 100 |
| K65 | AB924601 | 50 | 50 | 100 | 100 | 100 | NA | 100 | 100 | 100 |
| K66 | AB924602 | 33.33 | 50 | 100 | 100 | 100 | 100 | 100 | 100 | NA |
| K67 | AB924603 | 100 | 100 | 100 | 100 | 100 | NA | 100 | 100 | 100 |
| K68 | AB924604 | 100 | 100 | 100 | 100 | 100 | 100 | 100 | 100 | NA |
| K69 | AB924605 | 100 | 100 | 100 | 100 | 100 | NA | 100 | 100 | 100 |
| K70 | AB924606 | 25 | 100 | 100 | 100 | 100 | 100 | 100 | 100 | NA |
| K71 | AB924607 | 9.09 | 100 | 25 | 100 | 100 | NA | 100 | NA | 100 |
| K72 | AB924608 | 100 | 100 | 100 | 100 | 100 | NA | 100 | 100 | 100 |
| K74 | AB924609 | 25 | 33.33 | 100 | 100 | 100 | 100 | 100 | 100 | NA |
| K79 | AB924610 | 100 | 100 | 100 | 100 | 100 | 100 | 100 | 100 | NA |
| K80 | AB924611 | 11.11 | 100 | 100 | 100 | 100 | 100 | 100 | 100 | NA |
| K81 | AB924612 | 4 | 33.33 | 16.66 | 100 | 100 | 100 | 100 | 100 | NA |
| K82 | AB924613 | 100 | 100 | 100 | 100 | 100 | NA | 100 | 100 | 100 |
| KL103 | LT174574 | 4.16 | 100 | 100 | 100 | 100 | 100 | 100 | 100 | NA |
| KL105 | LT174575 | 2.56 | 100 | 100 | 100 | 100 | 100 | 100 | 100 | NA |
| KL106 | LT174576 | 100 | 100 | 33.33 | 100 | 100 | NA | 100 | 100 | 100 |
| KL108 | LT174577 | 100 | 100 | 100 | 100 | 100 | NA | 100 | NA | 100 |
| KL109 | LT174578 | 100 | 100 | 100 | 100 | 100 | 100 | 100 | NA | NA |
| KL110 | LT174579 | 4 | 33.33 | 50 | 100 | 100 | 100 | 100 | NA | NA |
| KL111 | LT174580 | 16.66 | 33.33 | 100 | 100 | 100 | NA | 100 | 100 | 100 |
| KL112 | LT174581 | 7.14 | 100 | 100 | 100 | 100 | 100 | 100 | 100 | NA |
| KL113 | LT174582 | 50 | 100 | 100 | 100 | 100 | NA | 100 | NA | 33.33 |
| KL114 | LT174583 | 20 | 100 | 100 | 100 | 100 | 100 | 100 | 100 | NA |
| KL115 | LT174584 | 4.76 | 50 | 100 | 100 | 100 | 100 | 100 | 100 | NA |
| KL116 | LT174585 | 9.09 | 100 | 50 | 100 | 100 | 100 | 100 | NA | NA |
| KL117 | LT174586 | 100 | 100 | 100 | 100 | 100 | 100 | 100 | 100 | NA |
| KL118 | LT174587 | 33.33 | 100 | 100 | 100 | 100 | 100 | 100 | NA | NA |
| KL119 | LT174588 | 100 | 100 | 100 | 100 | 100 | NA | 100 | 100 | 100 |
| KL120 | LT174589 | 2.77 | 100 | 16.66 | 100 | 100 | 100 | 100 | 100 | NA |
| KL121 | LT174590 | 16.66 | 100 | 100 | 100 | 100 | NA | 100 | 100 | 100 |
| KL122 | LT174591 | 50 | 100 | 50 | 100 | 100 | NA | 100 | 100 | 100 |
| KL123 | LT174592 | 100 | 100 | 100 | 100 | 100 | NA | 100 | 100 | 100 |
| KL124 | LT174593 | 4.76 | 100 | 100 | 100 | 100 | 100 | 100 | 100 | NA |
| KL125 | LT174594 | 4.16 | 100 | 100 | 100 | 100 | 100 | 100 | 100 | NA |
| KL126 | LT603702 | 50 | 100 | 50 | 100 | 100 | 100 | 100 | NA | NA |
| KL127 | LT603704 | 33.33 | 50 | 12.5 | 100 | 100 | 100 | NA | 100 | NA |
| KL130 | LT603706 | 3.84 | 100 | 100 | 100 | 100 | 100 | 100 | 100 | NA |
| KL131 | LT603707 | 6.66 | 100 | 14.28 | 100 | 100 | 100 | 100 | 100 | NA |

| | | | | | | | | | | |
|---|---|---|---|---|---|---|---|---|---|---|
| KL132 | LT603708 | 100 | 100 | 100 | 100 | 100 | NA | 100 | 100 | 100 |
| KL133 | LT603709 | 100 | 100 | 100 | 100 | 100 | NA | 100 | 100 | 100 |
| KL134 | LT603710 | 100 | 100 | 33.33 | 100 | 100 | ``````` | 100 | 100 | 100 |
| KL135 | LT603711 | 5.88 | 100 | 100 | 50 | 100 | 50 | 100 | 100 | NA |
| KL136 | LT603712 | 25 | 100 | 100 | 100 | 100 | NA | 100 | 100 | 100 |
| KL137 | LT603713 | 5 | 100 | 100 | 100 | 100 | 100 | 100 | 100 | NA |
| KL138 | LT603714 | 100 | 100 | 100 | 100 | 100 | 100 | 100 | 100 | NA |
| KL139 | LT603715 | 25 | 100 | 50 | 100 | 100 | NA | 100 | 100 | 100 |
| KL140 | LT603716 | 50 | 100 | 100 | 100 | 100 | NA | 100 | 100 | 100 |
| KL141 | LT603717 | 7.69 | 100 | 100 | 100 | 100 | 100 | 100 | 100 | NA |
| KL142 | LT603718 | 4.76 | 33.33 | 16.66 | 100 | 100 | NA | 100 | 100 | 100 |
| KL143 | LT603719 | 5 | 50 | 100 | 100 | 100 | 100 | 100 | 100 | NA |
| KL144 | LT603720 | 25 | 100 | 100 | 100 | 100 | NA | 100 | 100 | 100 |
| KL146 | LT603721 | 3.84 | 100 | 50 | 50 | 50 | 50 | 100 | 50 | NA |
| KL148 | LT603722 | 3.33 | 100 | 100 | 100 | 100 | NA | 100 | NA | 100 |
| KL149 | LT603723 | 6.66 | 50 | 100 | 100 | 100 | NA | 100 | NA | 100 |
| KL150 | KR007675 | 50 | 50 | 100 | 100 | 100 | 100 | 100 | NA | NA |
| KL153 | LT603725 | 100 | 100 | 100 | 100 | 100 | 100 | 100 | NA | NA |
| KL159 | LT603726 | 4.16 | 50 | 100 | 100 | 100 | NA | 100 | 100 | 100 |
| KN1 | AB924614 | 20 | 50 | 100 | 100 | 100 | 100 | 100 | 100 | NA |
| KN2 | LT174595 | 4 | 50 | 50 | 100 | 100 | 100 | 100 | 100 | NA |
| KN3 | LC189075 | 20 | 100 | 100 | 100 | 100 | NA | 100 | 100 | 100 |
| KL104 | Kaptiveref_KL 104 | 4 | 50 | 50 | 50 | 50 | NA | 100 | 100 | 50 |
| KL107 | Kaptiveref_KL 107 | 5.88 | 33.33 | 16.66 | 100 | 100 | 100 | NA | NA | NA |
| KL128 | Kaptiveref_KL 128 | 3.57 | 50 | 100 | 100 | 100 | 100 | 100 | 100 | NA |
| KL145 | Kaptiveref_Kl 145 | 100 | 100 | 100 | 100 | 100 | 100 | 100 | 100 | NA |
| KL147 | Kaptiveref_KL 147 | 4 | 50 | 100 | 100 | 100 | NA | 100 | NA | 100 |
| KL151 | Kaptiveref_KL 151 | 33.33 | 100 | 100 | 100 | 100 | NA | 100 | 100 | 100 |
| KL152 | Kaptiveref_KL 152 | 25 | 100 | 100 | 100 | 100 | NA | 100 | NA | 100 |
| KL154 | Kaptiveref_KL 154 | 4.16 | 100 | 50 | 50 | 50 | 50 | 50 | 50 | NA |
| KL155 | Kaptiveref_KL 155 | 25 | 50 | 50 | 100 | 100 | NA | 100 | 100 | 100 |
| KL157 | Kaptiveref_KL 157 | 100 | 100 | 100 | 100 | 100 | NA | 100 | NA | 100 |
| KL158 | Kaptiveref_KL 158 | 5 | 100 | 100 | 100 | 100 | NA | 100 | NA | 100 |
| KL160 | Kaptiveref_KL 160 | 50 | 50 | 100 | 100 | 100 | NA | 100 | 100 | 100 |

| | | | | | | | | | | |
|---|---|---|---|---|---|---|---|---|---|---|
| KL161 | Kaptiveref_KL 161 | 100 | 100 | 100 | 100 | 100 | NA | 100 | NA | 100 |
| KL162 | Kaptiveref_KL 162 | 3.03 | 100 | 100 | 100 | 100 | NA | 100 | 100 | 100 |
| KL163 | Kaptiveref_KL 163 | 6.25 | 50 | 50 | 50 | 50 | 100 | 100 | 50 | NA |
| KL164 | Kaptiveref_KL 164 | 100 | 100 | 100 | 100 | 100 | 100 | 100 | 100 | NA |
| KL165 | Kaptiveref_KL 165 | 25 | 100 | 50 | 100 | 100 | 100 | 100 | 100 | NA |
| AVERAGE RELIABILITY SCORE | | 37.13 | 80.83 | 79.88 | 94.28 | 96.43 | 97.18 | 98.54 | 97.52 | 93.81 |

**(B)**

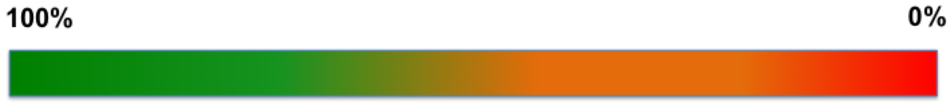


| Serotype | Genbank accession ID (Wzm) | Genbank accession ID (Wzt) | Reliability Score (%) | |
|---|---|---|---|---|
| | | | Wzm | Wzt |
| O1 | AAC98411.1 | AAC98413.1 | 33.3 | 33.3 |
| O2 | BAU36938.1 | BAU36939.1 | 33.3 | 33.3 |
| O2aeh | AVA30565.1 | AVA30566.1 | 100 | 100 |
| O2ac | BAU36940.1 | BAU36941.1 | 33.3 | 33.3 |
| O3 | AQZ41253.1 | AQZ41254.1 | 100 | 100 |
| O3 | BAU51051.1 | BAU51052.1 | 50 | 50 |
| O4 | ALX35080.1 | ALX35079.1 | 100 | 100 |
| O5 | BAN20050.1 | BAN20051.1 | 100 | 100 |
| O8 | AAC98405.1 | AAC98406.1 | 100 | 100 |
| O9 | BAU24812.1 | BAU24813.1 | 100 | 100 |
| O12 | BAN08506.1 | BAN08507.1 | 100 | 100 |
| OL101 | CZQ25251.1 | CZQ25252.1 | 100 | 100 |
| OL102 | CZQ25262.1 | CZQ25263.1 | 100 | 100 |
| OL103 | CZQ25266.1 | CZQ25267.1 | 100 | 100 |
| OL104 | CZQ25273.1 | CZQ25274.1 | 50 | 50 |
| Average reliability score | | | 83.3 | 83.3 |
