## Supplementary Table S6 for "K-PAM: A unified platform to distinguish *Klebsiella* species K- and O-antigen types, model antigen structures and identify hypervirulent strains"

**Table S6.** Test cases corresponding to clinically important *Klebsiella* species whose serotypes are undefined.

| Wzm Query ID | Wzt Query ID | Predicted O-type | | |
|---|---|---|---|---|
| | | Wzm | Wzt | Wzm & Wzt |
| AAF04380.1 | AAF04381.1 | O5 | O5 | O5 |
| KHF68821.1 | KHF68822.1 | O1, O2, O2ac | O1, O2, O2ac | O1, O2, O2ac |
| EXF40838.1 | EXF40837.1 | O3 | O3 | O3 |
| AAN06492.1 | AAN06493.1 | O12 | O12 | O12 |
| CCI88064.1 | CCI88065.1 | O1, O2, O2ac | O1, O2, O2ac | O1, O2, O2ac |
| CTQ06126.1 | CTQ06130.1 | O3 | O3 | O3 |
| PUH05602.1 | PUH05603.1 | O5 | O5 | O5 |
